# Structural basis of augment taurine uptake by taurine transporter alleviating cellular senescence

**DOI:** 10.1101/2024.11.13.623519

**Authors:** Heng Zhang, Nana Cui, Xiong Ma, H. Eric Xu

## Abstract

Taurine, a sulfur-containing amino acid, is critical for diverse physiological processes including liver function, cardiovascular health, and neurological development. Its deficiency has been linked to liver damage, cardiomyopathy, retinal degeneration, and accelerated aging. Despite its importance in metabolic regulation and disease prevention, the structural basis for taurine’s cellular uptake and its potential therapeutic applications have remained elusive. Here, we present high-resolution cryo-electron microscopy structures of the human taurine transporter (TauT) in multiple conformational states, revealing key insights into the taurine transport mechanism. We identify five distinct taurine-binding sites, elucidating a continuous pathway for taurine translocation across the cell membrane. Notably, we discover a novel gating mechanism involving a conserved salt bridge, which when disrupted, significantly enhances taurine uptake. Leveraging these structural insights, we demonstrate that augmented taurine uptake via TauT alleviates stress-induced senescence in biliary epithelial cells, potentially through the upregulation of the stress-responsive gene ATF3. Our findings not only advance the understanding of neurotransmitter transport but also unveil TauT as a promising therapeutic target for age-related liver diseases and other senescence-associated disorders. This work bridges structural biology with physiological function, offering new avenues for developing interventions to combat cellular aging and taurine-deficiency related diseases.

## Introduction

Taurine, a sulfur-containing amino acid, plays a crucial role in numerous physiological processes and is essential for the proper functioning of multiple organ systems^1^. This versatile molecule is involved in bile acid conjugation^2^, osmoregulation^3^, antioxidation^4,5^, and calcium homeostasis^6,7^. Taurine deficiency has been implicated in a wide array of health issues, including liver damage^8–10^, cardiomyopathy^11,12^, male reproductive impairments^13^, early embryonic developmental disorders^14,15^, reduced exercise capacity^16,17^, and accelerated aging^18^. The broad spectrum of taurine’s physiological functions underscores its critical importance in metabolic regulation and disease prevention across various tissues.

While endogenous taurine is synthesized through the cysteine sulfinic acid pathway^19–21^, dietary intake serves as the primary source of taurine in the body^22^. The absorption and cellular uptake of taurine are primarily facilitated by the taurine transporter (TauT)^23–25^, encoded by the *SLC6A6* gene. TauT belongs to the neurotransmitter sodium symporter family, specifically classified under the GABA transporters subfamily^26,27^. Given taurine’s vital role in cellular protection, the efficiency of TauT directly impacts the taurine shuttle and its protective effects on cells.

Recent studies have highlighted the severe consequences of impaired taurine transport, often resulting from genetic mutations or reduced TauT expression^9,12,28,29^. These consequences include tumor progression^30–32^, impaired liver ammonia detoxification^4,8,33^, retinal degeneration^28,29,34,35^, and cardiomyopathy^12,36^. Furthermore, emerging evidence suggests a strong link between TauT function and cellular senescence, a hallmark of aging^18^. Taurine uptake through TauT has been shown to mitigate stress-induced cellular aging, pointing to a potential therapeutic avenue for age-related diseases.

Despite significant advancements in structural understanding of related transporters such as human monoamine transporters^37–45^, GlyT1^46,47^, and GAT1^48–50^, fundamental questions regarding the transport mechanism of TauT remain unresolved. These include the specifics of ligand recognition and the conformational changes occurring between functional states. Elucidating these mechanisms is crucial for understanding how TauT efficiency impacts cellular aging and for developing novel therapeutic strategies to delay age-related decline and treat conditions like cirrhosis.

To address these critical knowledge gaps, our study presents high-resolution cryo-electron microscopy (cryo-EM) structures of TauT in multiple states: the apo state under both detergent and nanodisc conditions, and two distinct conformations when bound to taurine molecules. By combining these structural insights with functional assays, we provide a comprehensive understanding of the substrate recognition and transport mechanisms of TauT. Furthermore, we demonstrate that enhanced taurine uptake through TauT can alleviate cholangiocyte senescence, as evidenced by our experiments on doxorubicin-induced bile duct senescence. This work expands our understanding of TauT’s role in cellular protection and its potential therapeutic applications.

### Structural determination and architecture of human TauT

To elucidate the mechanisms of TauT-mediated taurine recognition and transport, we employed single-particle cryo-electron microscopy (cryo-EM) to determine the structure of human TauT in both apo and taurine-bound states. We expressed full-length wild-type TauT fused with an N-terminal Flag tag, without any fiducial marker.

Initially, we solubilized TauT in detergent (Dodecyl-β-D-maltoside, DDM) and obtained a 3.20 Å resolution map of TauT-DDM in the apo state (Extended Data Fig. 1a, d). However, this approach required a high protein concentration (22 mg/mL) to achieve homogeneous sample distribution (Extended Data Fig. 1c). To improve particle orientation distribution, we reconstituted purified TauT-DDM into nanodiscs using brain polar lipids and the scaffold protein Saposin A (Extended Data Fig. 1b). This strategy yielded diverse distribution of particle orientations, including top and bottom views, in contrast to the predominantly side views observed with TauT-DDM (Extended Data Fig. 1e).

Using this improved sample preparation, we determined high-resolution structures of TauT-Saposin A in multiple states: an apo state adopting an inward-open conformation at 3.02 Å resolution (Extended Data Fig. 1f), and taurine-bound states in both inward-open (2.75 Å) and occluded (3.30 Å) conformations (Extended Data Fig. 2). These density maps allowed us to fit nearly the entire polypeptide chain and accurately place most side chains.

The overall architecture of TauT conforms to the canonical LeuT fold^51^, a conserved structure among neurotransmitter transporters (Fig. 1a-b). Transmembrane helices TM1-5 are related to TM6-10 by a pseudo-twofold axis of symmetry, aligned approximately parallel to the membrane. TM1 and TM6 are each split into two shorter helices (TM1a/TM1b and TM6a/TM6b), which, along with adjacent transmembrane domains, sustain substrate transport. The extracellular surface of TauT is primarily composed of extracellular loops ECL2 (F153–K217), ECL3 (T266-D287), ECL4 (N355–P382), and ECL6 (K526-P539) (Fig. 1b).

**Fig.1.**
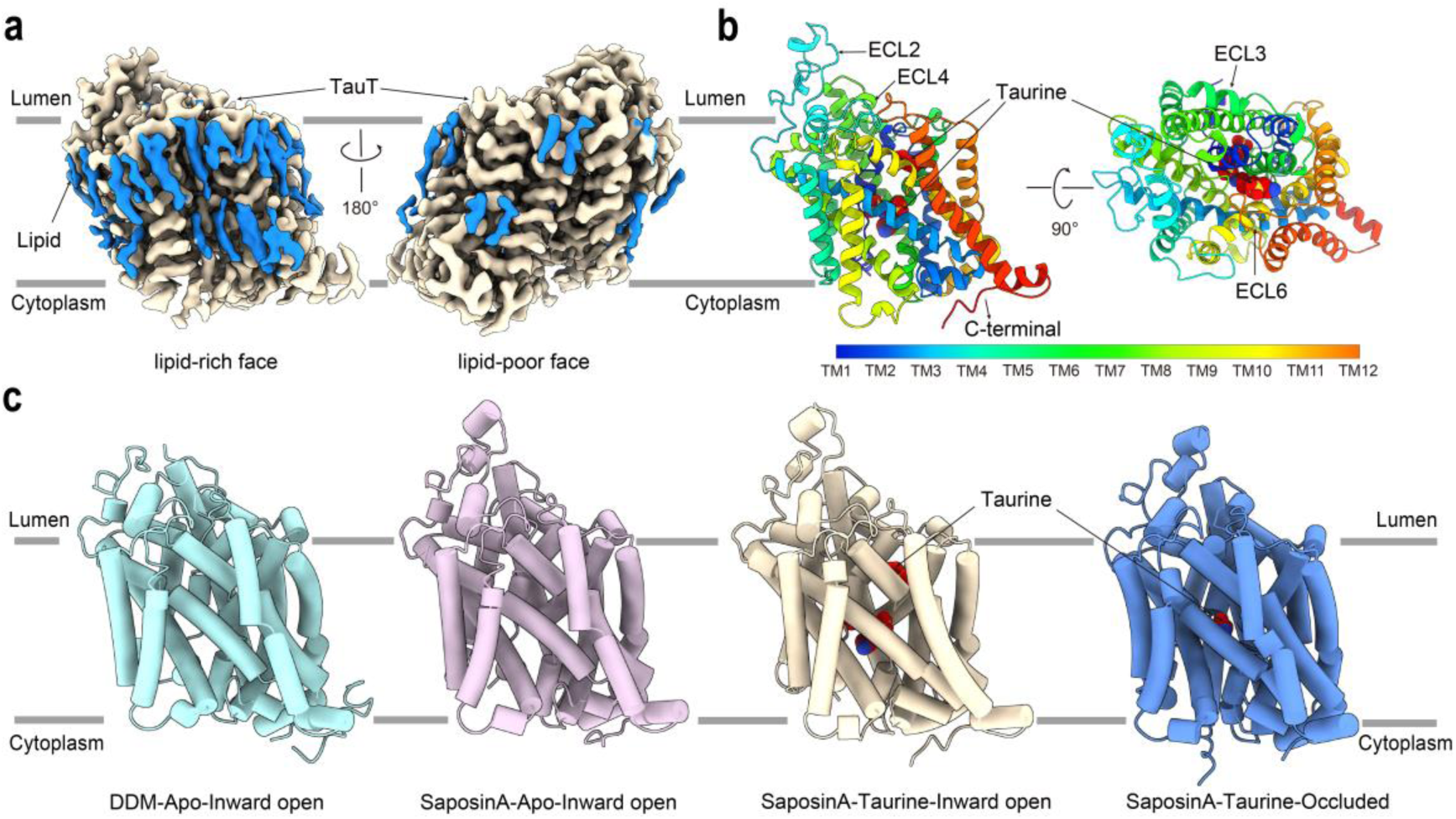
Architecture of the human taurine transporter TauT. **a**, the cryo-EM map of TauT bound to taurine reconstituted into SaposinA nanodisc. The lipid and TauT are colored in blue and gold, respectively. The map density contour level is 0.40. **b**, Side view and top view of the TauT bound taurine. The transmembrane helices are coloured as indicted and represented as cartoons. The taurine molecules are coloured in red and represented as spheres. **c**, the structures of Apo TauT in DDM and reconstituted into SaposinA nanodisc are coloured in pale turquoise and Thistle, respectively. the structures of taurine bound TauT reconstituted into SaposinA nanodisc adopting inward open conformation and occluded conformation are coloured in gold and blue, respectively.

Interestingly, we observed an asymmetric distribution of lipids surrounding TauT, similar to that seen in the norepinephrine transporter (NET)^41^ (Fig. 1a). Specifically, beyond the cholesterol molecule located at the classical cholesterol recognition site, we identified one cholesterol molecule in the cytoplasmic layer and three additional cholesterol molecules in the luminal layer based on the TauT-SaposinA density maps (Extended Data Fig. 2f). Despite the presence of lipid-rich regions overlapping with the dimeric interface seen in NET, TauT was consistently observed as a monomer in both detergent and nanodisc conditions (Fig. 1c).

Both the TauT-DDM and TauT-Saposin A structures in the apo state adopt an inward-open conformation, featuring a cytosol-accessible cavity through the intracellular vestibule (Fig. 1c). The two structures exhibit an overall root mean square deviation (RMSD) of 2.029 Å. Notably, TauT-DDM shows a 3.02 Å tilt of TM1a toward TM5 compared to TauT-Saposin A, with the most pronounced difference being the visibility of the C-terminus from A589 to S600 (Extended Data Fig. 3a).

Unlike the C-terminus of NET^39,41^, which forms multiple polar interactions with TM10 and TM11, the C-terminus of TauT remains flexible in both TauT-DDM and TauT-Saposin A (Extended Data Fig. 3b). This flexibility is intriguing, as previous studies have shown that phosphorylation of TauT’s C-terminus and intracellular domains is crucial for regulating its cell surface expression and transport activity^52,53^. The relatively flexible C-terminus in TauT, contrasting with the compact and stable C-termini observed in monoamine transporter structures, offers a structural template for future investigations into its role in modulating the cytoplasmic gate.

### Five taurine molecules bound in two conformations of TauT

To establish the structural basis of taurine transport by TauT, we incorporated 1 mM taurine throughout the purification process to ensure effective substrate binding. This strategy allowed us to capture TauT in two distinct conformational states, each providing insights into different stages of the transport cycle. Through iterative reconstitutions and meticulous local refinements of the cryo-EM data, we resolved the structures of these two complexes at resolutions of 2.75 Å and 3.30 Å, respectively (Extended Data Fig. 2a-c). The high-quality density maps enabled unambiguous assignment of taurine molecules and ions within the transporter (Extended Data Fig. 2d-g).

In both structures, the outward-facing entrance of TauT is closed, preventing extracellular access to the binding sites (Fig. 2a, d). However, the conformations differ markedly in the orientation of transmembrane helix TM1a and the number of bound taurine molecules. In the first structure, we observed a single taurine molecule bound at the central substrate-binding site. Here, TM1a is oriented almost vertically relative to the membrane plane, effectively sealing off the binding site from the cytoplasmic side. This configuration represents an occluded conformation (Fig. 2a), wherein the substrate is locked within the transporter and inaccessible from either side of the membrane—a state analogous to that observed in other neurotransmitter sodium symporters^41,44,47,48^(Extended Data Fig. 4c-d).

**Fig.2.**
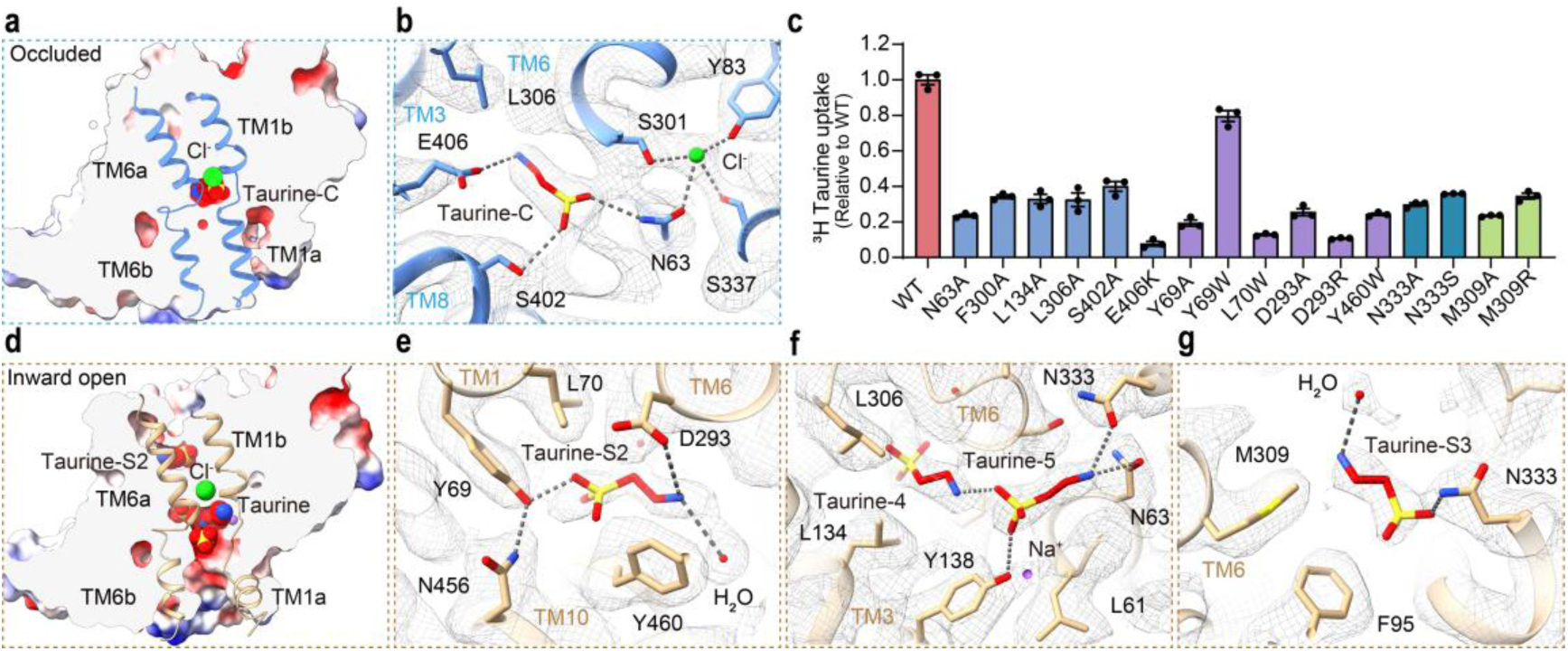
Illustration of taurine binding sites. **a**, Central taurine binding site of TauT in the occluded conformation. The taurine and chloride ion are coloured in red and green represented as spheres, respectively. **b**, Electron density and interactions of Central binding pocket of taurine and TauT. **c**, Comparative evaluation of the [^3^H] taurine transportation activity of TauT mutants in relation to the WT. Data are mean ± s.e.m. from three independent assays, each with one measurement (n = 3). The transportation rates of all mutations are significantly lower than that of wild-type TauT. P < 0.0001 for all mutants, one-way ANOVA test. **d**, Four taurine molecules bound of TauT in the inward open conformation. The taurine molecules and chloride ion are coloured in red and green represented as spheres, respectively. **e**, Electron density and interactions of taurine-S2 binding pocket of taurine and TauT. **f**, Electron density and interactions of taurine-4 and taurine-5 binding pockets of taurine and TauT. **g**, Electron density and interactions of taurine-S3 binding pocket of taurine and TauT.

In contrast, the second structure revealed four taurine molecules bound at distinct sites along the translocation pathway. In this conformation, TM1a tilts upward toward the membrane plane, creating an open pathway to the cytoplasmic side (Fig. 2d). This movement opens the cytosolic entrance, resembling the inward-open conformation observed in the apo state of TauT. The presence of multiple taurine molecules suggests the existence of additional substrate-binding sites, providing new insights into the stepwise mechanism of taurine transport through TauT.

To confirm that these binding sites are specific to the taurine-bound states, we compared them with the apo-state structures of TauT obtained under both detergent and nanodisc conditions (Extended Data Fig. 4a-b). In the apo-state structures, no clear density corresponding to taurine was observed at the equivalent sites. This absence underscores that the observed taurine densities in our complexes are due to specific substrate binding rather than nonspecific interactions or artifacts of sample preparation.

### Central binding site of Taurine in the occluded conformation of TauT

In the occluded conformation, we observed a single taurine molecule bound in the central binding site, with its long axis oriented nearly parallel to the membrane plane (Fig. 2a-b). The central binding site can be divided into three subsites (A, B, and C), mirroring the arrangement reported in monoamine transporters. At subsite A, the sulfonic acid group of taurine is nestled in the hinge region between TM1 and TM6. This group forms hydrogen bonds with the main chain atoms of F58^TM1^, G60^TM1^, G62^TM1^, and S301^TM6^, while also establishing salt bridges with the side chains of N63^TM1^ and S402^TM8^ (Fig. 2b). The aminoethane group of taurine interacts with subsites B and C. In subsite B, it forms a salt bridge with E406^TM8^ and engages in hydrophobic interactions with L134^TM3^ and L306^TM6^. Subsite C contributes an ion-π interaction between the amino group and F300^TM6^ (Fig. 2b).

To validate the functional importance of these interactions, we performed site-directed mutagenesis and [^3^H]taurine uptake assays. Mutations at all three subsites resulted in markedly decreased transport activity of TauT, with residues N63 and E406 showing particularly pronounced effects (Fig. 2c). Importantly, flow cytometry analysis revealed no significant differences in surface expression between wild-type TauT and these mutants, confirming that the observed effects were due to impaired transport rather than trafficking defects (Extended Data Fig. 6).

While the overall architecture of the central binding site is conserved across neurotransmitter sodium transporters, subtle differences in amino acid distribution can lead to variations in substrate binding poses. Comparing our occluded TauT structure with other transporters revealed interesting distinctions (Extended Data Fig. 4c-d). The taurine binding site in TauT closely overlaps with that of GABA in GAT1^48^, but the presence of E406 in TauT causes the plane of taurine’s amino group to tilt by approximately 24° relative to the carboxyl group of GABA (Extended Data Fig. 4c).

Notably, the binding orientation of taurine in TauT is reversed compared to monoamines like NE^41^ in their respective transporters. In monoamine transporters, the primary amine is typically anchored in subsite A, with the electronegative group accommodated by subsites B and C. In contrast, taurine’s sulfonic acid group occupies subsite A in TauT. This reversal in orientation can be attributed to the different locations of key residues, such as E406 in TauT compared to D75 in NET (Fig. 2b, Extended Data Fig. 4d). Together, these structural insights, coupled with our functional data, provide a detailed molecular understanding of taurine recognition by TauT and highlight the subtle adaptations that allow this transporter to specifically accommodate its unique substrate.

### Allosteric binding sites of Taurine molecules in the Inward-open conformation of TauT

While substrate allosteric binding sites are well-established in monoamine transporters^38,39^, such sites have not been previously validated for other SLC6 family members. In our study, we discovered four additional taurine-binding sites in the inward-open conformation of TauT, in addition to the single taurine molecule bound in the occluded state (Fig. 2d-g). These four taurine molecules are sequentially aligned along the translocation pathway from the outward to the inward entrance, despite the outward permeation pathway being closed.

The first taurine molecule, referred to as taurine-S2, is trapped in the outward entrance, located above the central binding site and slightly lower than the S2 site observed in the serotonin transporter (SERT)^38^ (Extended Data Fig. 4e). Well-resolved water molecules in this region facilitate hydrogen bond interactions between the aminoethane group of taurine and the side chains of D293^TM6^ and Y460^TM10^ (Fig. 2e). Additionally, the sulfonic acid group of taurine forms a hydrogen bond with the phenolic hydroxyl group of Y69^TM1^, which is stabilized by N456^TM10^. The residues involved in taurine binding at the S2 site, including L70^TM1^, are highly conserved within the GABA transporter subfamily of SLC6 transporters. Functional assays demonstrated that mutations in the S2 binding pocket significantly reduce the transport activity of TauT (Fig. 2c), highlighting the critical role of this site in substrate recognition.

In contrast to SERT, where conserved amino acids like D328^TM6^ and S559^TM11^ form the allosteric binding pocket for 5-HT, TauT possesses different residues at analogous positions. In SERT, E494^TM10^ and Y579^TM12^ create a salt bridge and a T-shaped π–π stacking interaction with 5-HT, respectively. In TauT, these positions are occupied by Y460^TM10^ and W547^TM12^, which reduce the available space and prevent taurine from binding at this site (Extended Data Fig. 4e). Moreover, the negatively charged side chain of E494^TM10^ and an approximate 1.5 Å tilt of TM10 toward TM11 introduce significant hindrance to taurine binding. Additional steric obstruction arises from tilts of approximately 2.4 Å in extracellular loop 3 (ECL3) and 2.2 Å in ECL4 toward the NE-S2 site^39^ in the norepinephrine transporter (NET), along with the bulky side chain of F276^ECL3^ (Extended Data Fig. 4e).

At the central site, we observed two taurine molecules, termed taurine-4 and taurine-5, arranged in a head-to-tail configuration (Fig. 2f). Compared to the binding of taurine in the occluded state of TauT and GABA in GAT1^48^, these two taurine molecules exhibit flips of approximately 180° and 160°, respectively (Extended Data Fig. 5a). This significant change in binding pose is associated with conformational shifts within TauT; specifically, TM1a rotates by about 28° toward the membrane plane, and TM8 shifts approximately 3 Å away from the central site in the inward-open conformation (Extended Data Fig. 5a-b). These coordinated movements create a larger cavity, providing taurine with increased opportunities to enter the inward-facing permeation pathway and access the cytoplasm.

Another taurine molecule, referred to as taurine-S3, was identified near the central inward-facing entrance (Fig. 2g). The sulfonic acid group of taurine-S3 forms a polar interaction with N333^TM7^, while the amino group is stabilized through coordination with M309^TM6^ via water molecules. Mutation of M309 to either alanine (with a shorter side chain) or arginine (with a basic side chain) significantly impaired taurine transport capacity (Fig. 2c).

The structural insights gained from the taurine-bound TauT in both the occluded and inward-open conformations provide a valuable framework for understanding the mechanics of substrate recognition and translocation. The identification of multiple taurine-binding sites in the inward-open state not only expands our comprehension of TauT’s transport mechanism but also suggests a broader applicability of allosteric binding in other neurotransmitter sodium transporters.

### Δ464S TauT exhibits augment of taurine uptake

When we superimposed the five taurine molecules found in different structural states of TauT, a continuous passage from the lumen-facing permeation pathway to the cytoplasm-facing permeation pathway emerged (Fig. 3a). The presence of taurine-S2, trapped in the outward entrance of an inward-open structure (Fig. 3b), was particularly intriguing as no previous experimental data had indicated neurotransmitter binding at this site in an inward-open transporter.

**Fig.3.**
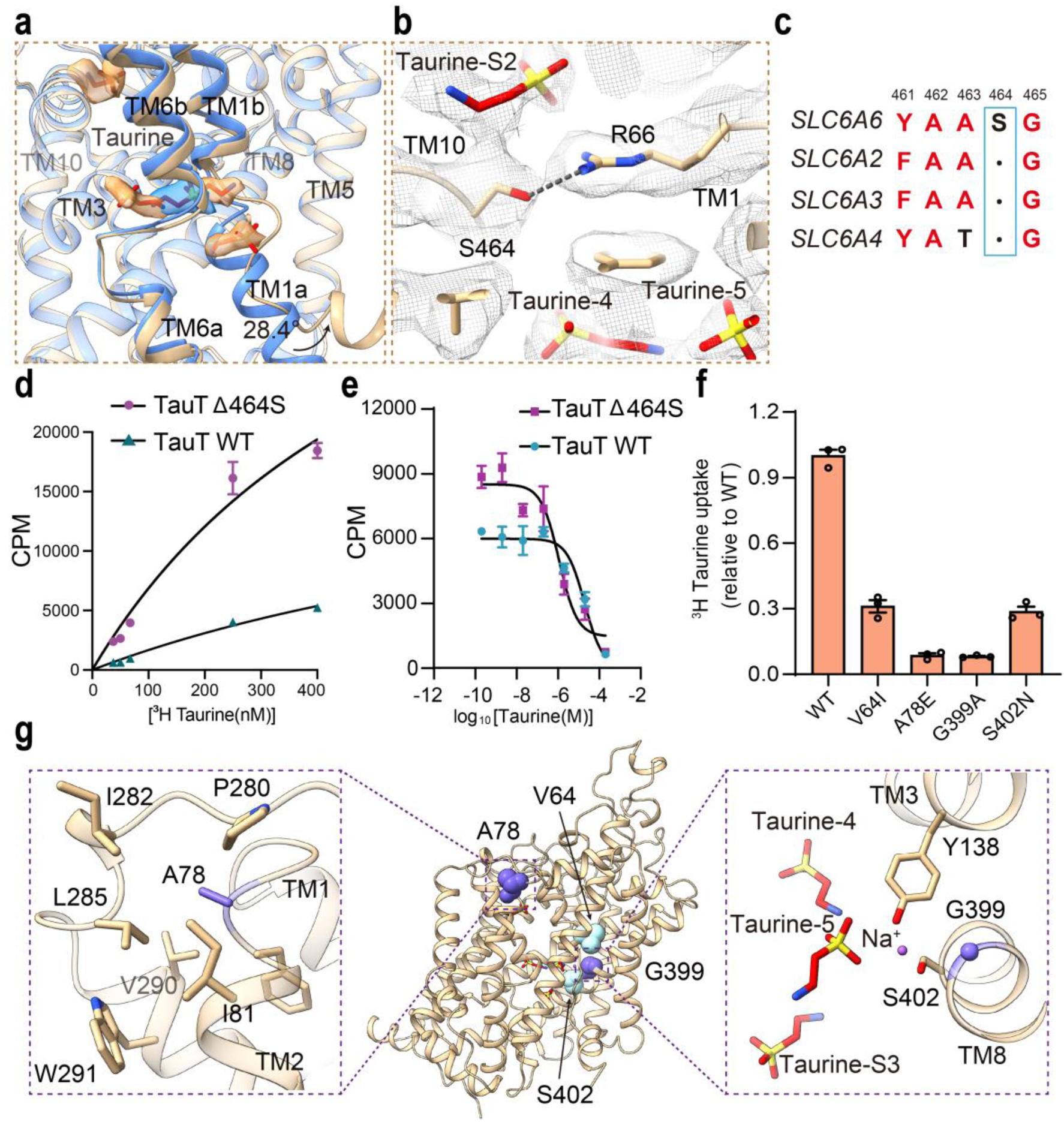
Mutations affect the transport capacity of TauT. **a**, Superstition of taurine molecules in the occluded and inward-open conformations. **b**, Electron density and interactions of outward entrance gate of TauT. **c**, Sequence alignment of TM8 of monoamine transporters and GABA subtype transporters, only the amino acids near the S464 shown. **d**, Concentration-dependence of [^3^H]taurine uptake by HEK293 cells expressing wild-type TauT (blue) and mutant Δ464S (red), respectively. Km = 1.12 μM for wild-type TauT, Km = 0.62 μM for mutant Δ464S, Data are shown as mean ± SD; n = 3 biological replicates. **e**, Uptake of [^3^H]taurine into HEK293 cells expressing wild-type TauT (blue) and mutant Δ464 (red) in the presence of increasing concentrations of unlabeled taurine. pIC50 for wild-type TauT is 4.70 ± 0.23, and pIC50 for mutant Δ464S is 5.64 ± 0. 45. **f**, Comparative evaluation of the [^3^H] taurine transportation activity of mutants related to reducing the transport capacity of TauT in relation to the WT. Data are mean ± s.e.m. from three independent assays, each with one measurement (n = 3). The transportation rates of all mutations are significantly lower than that of wild-type TauT. P < 0.0001 for all mutants, one-way ANOVA test. **g**, These positions of the mutations related to reducing the transport capacity of TauT mapped into inward-open conformation and taurine-bound TauT.

Previous structure-based predictions^27^ suggested that the gating of TauT’s outward permeation pathway involves two highly conserved amino acids, R66 and D459. In our structure, we found that R66, located under the taurine-S2, not only forms a salt bridge with D459 but also with S464 (Extended Data Fig. 7a-c). Further investigation through sequence alignment revealed that monoamine transporters lack a residue corresponding to S464 (Fig. 3c). These observations led us to hypothesize that the additional salt bridge between R66 and S464 in TauT impedes substrate access to the central binding site from the outward entrance.

To test this hypothesis, we engineered a Δ464S mutation in TauT. Transport experiments yielded striking results: compared to the wild type, the Δ464S mutant exhibited significantly enhanced substrate transport capacity (Fig. 3d). Substrate competition uptake assays provided further evidence, revealing a pIC50 value of 5.6 for Δ464S compared to 4.7 for WT, indicating both higher affinity and greater transport capacity for taurine in the mutant (Fig. 3e). These findings strongly suggest that disrupting this additional gate substantially enhances TauT’s substrate transport capability.

This enhancement is particularly significant in contrast to monoamine transporters^37–44^, GAT1^46,47^ and GlyT1^48–50^, which are often targeted by inhibitors in clinical applications. In contrast, TauT-related diseases are associated with reduced transport function. Clinical mutations^29^, such as A78E and G399A, linked to hypotaurinemic retinal degeneration and cardiomyopathy, respectively, show significantly reduced taurine transport activity compared to wild-type TauT (Fig. 3f).

The A78E mutation, located in the hinge region between TM1b and TM2, disrupts the hydrophobic environment essential for the coordinated movement of TM1b and TM2, which is crucial for TauT’s outward-to-inward conformational transition. Similarly, the G399A mutation impairs sodium ion binding, which is vital for taurine transport driven by sodium and chloride ions (Fig. 3g). Additionally, other loss-of-function mutations, such as V64I and S402N, also diminish TauT’s substrate transport capacity by affecting sodium ion binding sites (Fig. 3f-g). Thus, enhancing TauT’s transport capacity, such as through the Δ464S mutation, represents a promising therapeutic approach to mitigate TauT-related diseases characterized by reduced taurine uptake.

### Augmented Taurine Uptake by Taut alleviating biliary epithelial cell aging

Taurine plays a crucial role in liver function, essential for bile acid metabolism and maintaining healthy liver and biliary systems^2^. To investigate TauT’s role in the liver, we utilized in situ multi-color immunofluorescence to visualize TauT expression, revealing that TauT is predominantly localized in CK7-positive biliary epithelial cells (Fig. 4a). Notably, TauT expression was significantly higher in healthy liver tissues compared to those affected by cirrhosis (Fig. 4b-d). This observed decline in TauT expression in cirrhotic tissues suggests a potential link between reduced taurine transport and impaired liver function, underscoring the importance of TauT in maintaining hepatic and biliary homeostasis.

**Fig.4.**
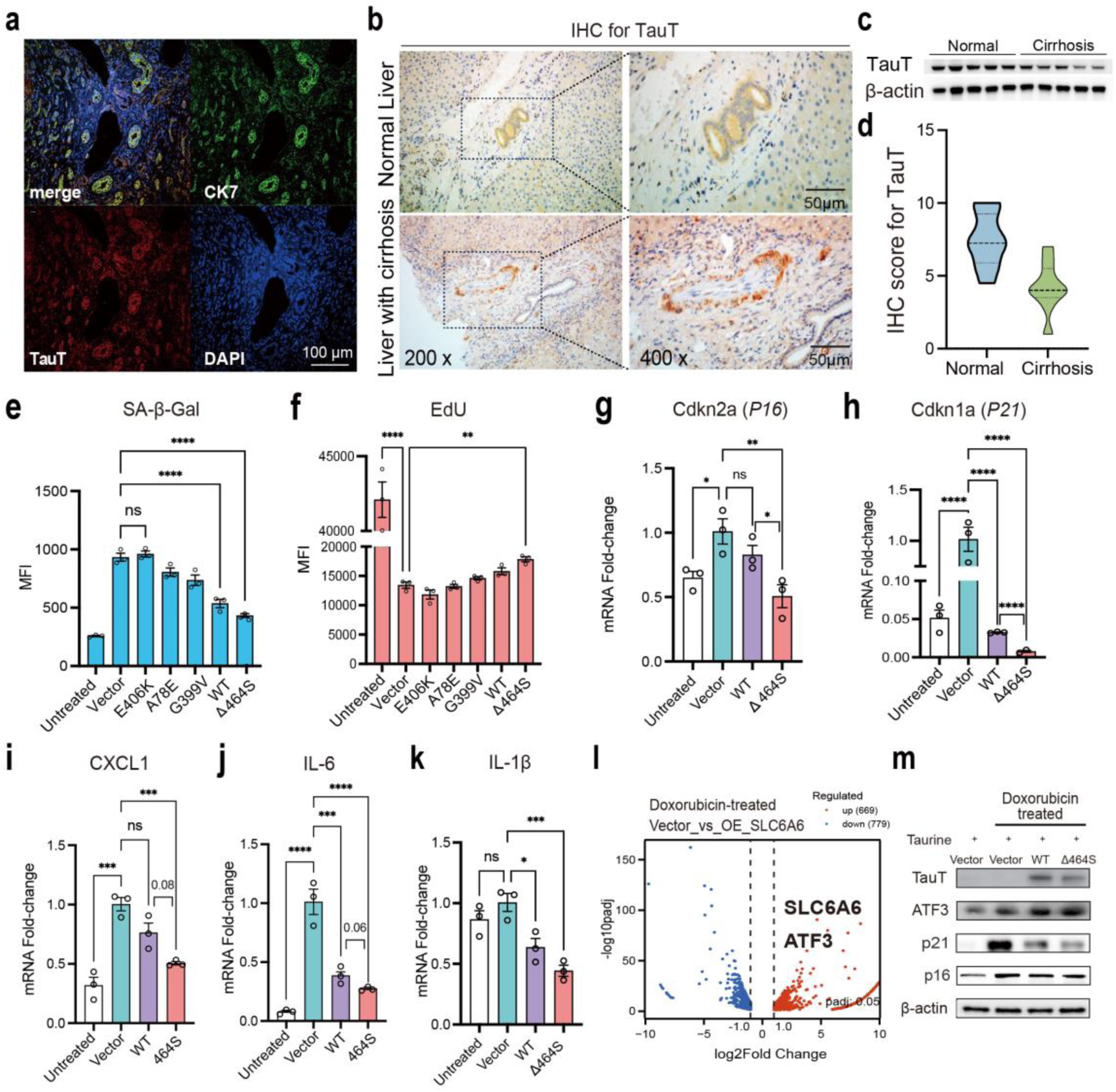
Taurine transported by TauT into biliary epithelial cells regulates classical hallmarks of aging. **a**, Representative TauT and CK7 staining in bile duct tissues from normal liver. Scale bar, 100 μm. **b**, Representative staining of TauT in normal liver and cirrhosis liver. Scale bar, 50 μm. **c**, TauT protein expression levels in normal livers and cirrhosis livers. **d**, Immunohistochemistry evaluations of TauT expression in normal livers and cirrhosis livers. **e**, MFI quantifications of SA β-galactosidase expression in doxorubicin treated and transfected with TauT or TauT mutants of RBE cells. **f**, MFI quantifications of the proliferative cells in doxorubicin treated and transfected with TauT or TauT mutants of RBE cells. **g-h**, qPCR showing the relative expression of *P16* (**g**) and *P21* (**h**) in doxorubicin treated and transfected with wild type TauT or TauT-Δ464S of RBE cells. **i-k,** qPCR showing the relative expression of SASPs including CXCL1 (**i**), IL-6 (**j**), and IL-1β (**k**) in doxorubicin treated and transfected with wild type TauT or TauT-Δ464S of RBE cells. Data are mean ± s.e.m. from three independent assays, each with one measurement (n = 3). Statistical analysis was performed with Graph Pad Prism 10 using one-way ANOVA test. All values are means ± SEM. p ≤ 0.0001****,p ≤ 0.001***, p ≤ 0.01**, p ≤ 0.05*, p > 0.05ns are versus WT or control, all. Unlabeled groups represent no statistical difference. **l**, Volcano map of RBE cells transfected TauT (*SLC6A6*) versus transfected vector. **m**, Western blot of TauT, ATDF3, *P16* and *P21* in doxorubicin treated and transfected with wild type TauT or TauT-Δ464S of RBE cells.

To determine whether the reduction in TauT expression contributes to bile duct aging, we treated RBE cells (epithelial neoplasm cells of bile duct) with Doxorubicin (Dox)^54^, a chemotherapeutic agent known to induce senescence and elevate senescence-associated secretory phenotype (SASP) markers. In RBE cells transfected with TauT mutants A78E, G399V, and E406K, Dox-induced senescence-associated β-galactosidase (SA-β-gal) staining showed no significant change compared to control cells (Fig. 4e, Extended Data Fig. 8a-b). In contrast, cells transfected with wild-type TauT and the Δ464S mutant exhibited a marked reduction in SA-β-gal staining, indicating reduced senescence. Similarly, GLB1 mRNA levels, a marker of senescence, were significantly decreased in cells expressing wild-type TauT and even more so in cells expressing TauT-Δ464S (Extended Data Fig. 8d).

Furthermore, wild-type TauT and TauT-Δ464S partially restored the proliferation deficiency induced by Dox, with the Δ464S mutant demonstrating greater improvement than wild-type TauT (Fig. 4f, Extended Data Fig. 8c, e). Conversely, TauT mutants A78E, G399V, and E406K did not show significant changes in cell proliferation. Analysis of SASP markers revealed that transfection with wild-type TauT and TauT-Δ464S significantly rescued the Dox-induced aberrant expression of genes such as p16, p21, CXCL1, IL-6, and IL-1β (Fig. 4g-k). These results indicate that enhanced taurine uptake via TauT mitigates stress-induced cellular aging and inflammation in biliary epithelial cells.

Consistent with the Δ464 mutation’s ability to enhance taurine transport activity, TauT-Δ464S exhibited superior performance in mitigating Dox-induced senescence of RBE cells compared to wild-type TauT (Fig. 4g–i). This suggests that increased taurine uptake through TauT confers protective effects against cellular aging. To elucidate the underlying mechanisms, we performed RNA sequencing (RNA-seq) on RBE cells transfected with a control vector and wild-type TauT, identifying 2,065 differentially expressed genes (DEGs; fold change >1.5, p < 0.05) (Fig. 4l). Kyoto Encyclopedia of Genes and Genomes (KEGG) pathway enrichment analysis revealed that genes related to the “nucleus” were significantly enriched following TauT overexpression (Extended Data Fig. 9).

Among these DEGs, Activating Transcription Factor 3 (ATF3), an early stress-responsive gene, was significantly upregulated (Fig. 4l-m). ATF3 is critical for maintaining cellular homeostasis by coordinating serine and nucleotide metabolism ^55,56^, suppressing STING1^56,57^ (a key regulator of innate immune responses), and repressing apoptosis-related and immunity-related genes^58,59^. We validated the Dox-induced upregulation of ATF3 in RBE cells, which was further enhanced by transfection with wild-type TauT. Notably, transfection with TauT-Δ464S resulted in an even greater increase in ATF3 expression (Fig. 4m). These findings suggest that augmenting taurine uptake through wild-type TauT and TauT-Δ464S regulates ATF3 expression, thereby providing protective effects against stress-induced cellular aging in biliary epithelial cells.

## Discussion

Previous studies have suggested a key role for taurine transported by TauT in the arresting the aging process^18^, antitumor^30–32^, male reproduction and early embryo development^13–15^, detoxification^4,8,33^, exercise^16,17^, and protection of myocardial function^11,12^. Despite its broad range of functions, the structural basis for TauT’s transport activity and its interaction with taurine has remained elusive. Our study provides critical insights into these mechanisms, revealing five distinct taurine-binding sites that form a continuous permeation pathway from the lumen-facing to the cytoplasm-facing regions in both occluded and inward-open conformations. And, four taurine molecules were bound together in an inwardly open conformation, which may have resulted from signal superposition during data processing, where taurine molecule signals were superimposed at multiple sites of the same conformational structure.

By integrating our findings with AlphaFold-predicted^60^ TauT structure, we have mapped the conformational transitions that TauT undergoes during the transport cycle (Extended Data Fig. 7d). This comprehensive structural view unveils different substrate-binding modes across various conformations, significantly advancing our understanding of TauT’s transport mechanism. Structural analysis highlighted the pivotal roles of discontinuous helices TM1 and TM6 in substrate recognition and conformational changes, aligning with observations in other neurotransmitter transporters and underscoring the conserved mechanism of the LeuT-fold transporters (Extended Data Fig. 7d). Notably, we identified a conserved salt bridge as a key gating mechanism, where its disruption significantly enhances taurine uptake, revealing a novel regulatory feature with potential implications for transporter modulation and therapeutic targeting.

Recent articles have emphasized taurine’s crucial role in anti-aging^18^, weight loss^61^, and other health benefits^1^. Additionally, studies have suggested that taurine uptake via TauT may facilitate tumor cell immune evasion, while taurine supplementation has been shown to rejuvenate exhausted CD8^+^ T cells, thereby enhancing the efficacy of cancer therapies. In light of these findings, developing strategies to deliver taurine to specific tissues to improve stress resilience represents a promising direction. Our research indicates that both wild-type TauT and the Δ464S mutant (disrupting the R66-S464 salt bridge and significantly increases taurine transport) can upregulate the early stress-responsive gene ATF3, thereby reducing senescence in bile duct epithelial cells. Notably, the Δ464S mutation offered greater protection against stress-induced aging compared to wild-type TauT, pointing to a potential therapeutic strategy for enhancing taurine uptake in pathological conditions.

## Methods

### Expression and Purification of TauT

The coding sequences for human wild-type TauT (SLC6A6) were cloned into the pEG BacMam vector. A 1xFlag tag was placed at the N-terminus of the protein, connected by a GGS linker. TauT was expressed in HEK293S GnTI− cells cultured in SMM-293TII medium (Cat#: M293TII, SinoBiology) supplemented with 2% fetal bovine serum (Gibco). Cells were harvested by centrifugation (1,000 × g, 10 min, 4 °C) and lysed using a Dounce tissue grinder (Merck Millipore) in buffer (50 mM HEPES pH 7.5, 150 mM NaCl) supplemented with 10 µg/mL aprotinin (HY-P0017, MedChemExpress), 4 µg/mL leupeptin (HY-18234, MedChemExpress), and 3 µg/mL pepstatin A (HY-P0018, MedChemExpress). Cell debris was removed by centrifugation (8,000 × g, 25 min, 4 °C), and the membrane fraction was collected via ultracentrifugation (100,000 × g, 1 h, 4 °C). The membrane pellet was solubilized in buffer (50 mM HEPES pH 7.5, 150 mM NaCl, 1% (w/v) n-dodecyl-β-D-maltoside (DDM, Anatrace), 0.1% (w/v) cholesteryl hemisuccinate (CHS, Anatrace)) for 2.5 h at 4 °C. The insoluble material was removed by centrifugation (30,000 × g, 1 h, 4 °C). The supernatant was incubated with anti-DYKDDDDK G1 affinity resin (GenScript) for 2 h at 4 °C, and the bound protein was eluted with 0.2 mg/mL DYKDDDDK peptide (GenScript). TauT was further purified via size-exclusion chromatography (SEC) on a Superdex 200 Increase 10/300 GL column (GE Healthcare), equilibrated with SEC buffer (20 mM HEPES pH 7.5, 150 mM NaCl, 0.012% DDM). Peak fractions were collected and concentrated to 22.0 mg/mL using a 100 kDa MWCO Amicon centrifugal filter (Merck Millipore).

### Nanodisc-TauT Reconstitution

Brain polar lipids (Avanti Polar Lipid) were dissolved in chloroform, dried under argon gas to form a lipid film, and stored under vacuum overnight. The lipid film was resuspended at 150 mM in buffer containing 20 mM Tris, pH 7.5, 100 mM NaCl, and 0.2% DDM. TauT, SaposinA (membrane scaffold protein), and lipids were mixed at a molar ratio of 1:10:20 in buffer containing 20 mM HEPES pH 7.5, 100 mM NaCl, and 10 mM sodium cholate. The mixture was incubated for 2 h at 4 °C. Nanodisc formation was initiated by adding 320 mg/mL equilibrated Bio-Beads SM-2 resin (Bio-Rad), followed by constant agitation overnight at 4 °C. The sample was purified using SEC on a Superdex 200 Increase 10/300 column (GE Healthcare) equilibrated with SEC buffer (20 mM HEPES pH 7.5, 150 mM NaCl). SEC peak fractions were pooled and concentrated to 1.2 mg/mL for cryo-EM sample preparation.

### Grid Preparation and Data Acquisition

For the cryo-EM structure determination of TauT in the apo and taurine-bound states (1 mM taurine, HY-B0351, MedChemExpress), 2.5 μL of purified protein was applied to glow-discharged (SuPro Instruments) holey carbon grids (Quantifoil R1.2/1.3 Au 300 mesh). Excess liquid was blotted off in a Vitrobot Mark IV (Thermo Fisher Scientific) at 8°C with 100% humidity for 4.5 seconds (blotting force of 1). The grids were flash-frozen in liquid ethane. Cryo-EM imaging was performed using a 300 kV Titan Krios G4 (FEI) equipped with a cold field emission gun (E-CFEG) and Falcon 4i direct electron detector. Data were collected at a pixel size of 0.73 Å, with a total dose of 50 e/Å² over 36 frames, at the Shanghai Advanced Center for Electron Microscopy. Motion correction was carried out using MotionCor2.

### Data Processing

For the TauT-DDM-Apo dataset, 9,243 dose-weighted micrographs were imported into cryoSPARC v4.3.1, and contrast transfer function (CTF) parameters were calculated using Patch CTF. Micrographs with CTF resolution worse than 3.8 Å were discarded. Particles were initially selected using a blob picker (100 Å diameter). After several rounds of 2D classification, 117,305 high-quality particles were retained for 3D reconstruction and refinement. A 3.86 Å class was selected as a good reference and used as a seed for further template picking. After multiple rounds of seed-facilitated multi-reference 3D classification, 129,662 particles were refined to yield a final map of TauT at 3.20 Å resolution as a monomer (Extended Data Fig. 1). Similar workflow was applied to the TauT-SaposinA datasets, resolving TauT maps at 3.02 Å for the apo state and at 2.75 Å and 3.30 Å for the taurine-bound state (Extended Data Fig. 2).

### Model Building and Refinement

The initial model of TauT was generated using AlphaFold2. The model was then fit into the cryo-EM density map using UCSF ChimeraX. Manual adjustments were made iteratively in COOT to match the density more closely. Ligand restraint files were generated with PHENIX 1.17_elbow, and models were refined against the cryo-EM maps using phenix.real_space_refine.

### Cell culture

We took advantage of human ICC cell line (RBE) and as in vitro models to explore the mechanism of TauT in transport assay and aging assay. RBE and HEK293T cell lines were purchased from National Collection of Authenticated Cell Cultures (Shanghai, China). RBE cells were cultured in RPMI Medium 1640 supplemented with 10% fetal bovine serum and 1% penicillin/streptomycin. 293T cells were cultured in Dulbecco’s modified Eagle medium supplemented with 10% fetal bovine serum and penicillin/streptomycin. Cells were cultured in a humidified chamber with 5% CO_2_ at 37°C.

### [^3^H]Taurine Uptake Assay

HEK293T cells were transiently transfected with wild-type or mutant TauT and seeded into 6-well plates at a density of 20,000 cells/well in 2 mL of media, then incubated at 37 °C with 5% CO_2_ for 24 hours. The cells were detached with 0.25% trypsin and reseeded into 96-well plates at 10,000 cells/well overnight. Cells were washed twice and incubated with blocking buffer (DMEM supplemented with 25 mM HEPES and 0.1% BSA, pH 7.4) for 2 hours at 37 °C. For competition transport assays, radiolabeled [^3^H]Taurine (6 nM) and unlabeled competitor (20 pM to 200 μM) were added to cells in blocking buffer and incubated at room temperature for 3 hours. After incubation, cells were washed three times with ice-cold PBS and lysed in 50 μL of lysis buffer (PBS with 20 mM Tris-HCl and 1% Triton X-100, pH 7.4). Radioactivity was measured (counts per minute, CPM) using a scintillation counter (MicroBeta 2 Plate Counter) with a scintillation cocktail (OptiPhase SuperMix, PerkinElmer). Data were analyzed via nonlinear regression in GraphPad Prism v10.

### Detection of surface expression of TauT

HEK293T cells were seeded in 96-well plates as described above and then transiently transfected with wild-type or mutations of TauT. The plasmid is consistence with the structural determination. The DNA–Lipofectamine 3000 transfection mixture containing 1 μg plasmid DNA and 2 μl Lipofectamine 3000 (Invitrogen) in Opti-MEM medium (Gibco) was applied. After 24 h transfection, the collected cells were suspended in PBS and washed with PBS containing 1% (w/v) BSA, followed by blocking with PBS containing 10% bovine serum (Sangon Biotech) at 4 °C for 1 h. Cells were then incubated with primary anti-Flag-M2-FITC antibody (Sigma) diluted with PBS containing 2% (w/v) BSA and 0.06% Triton X100 at a ratio of 1:100 or the same concentration of rabbit monoclonal IgG (isotype control, Abcam) for 10 min at 4 °. Fluorescence intensity was quantified using LSDFortessa X20 (BD). Approximately 10,000 cell events were collected per sample, and data were normalized to the wild-type TauT.

### Human subjects and samples

We compare the total protein levels in healthy (n = 5) and cirrhotic (n = 5) human liver tissues by conducting Western blot analysis. Demographics and characteristics of patients are presented at Extended Data Table 2. Histology of IHC-stained and IF-stained sections were observed using light microscopy and laser confocal microscopy, respectively. Three to five random fields were chosen for each section, and analyzed by two “blinded” hepatic pathologists. All individuals provided written informed consent. The study was carried out under the principles of the Declaration of Helsinki and approved by the Ethics Committee of Renji Hospital, Shanghai Jiao Tong University.

### Immunohistochemistry and immunofluorescence in human liver tissue

Immunohistochemical staining of TauT was performed by Leica Bond system (Leica, Germany). Paraffin-embedded liver tissues were treated with heat mediated antigen retrieval using citrate buffer (pH 6) for 20 min followed by incubation with an anti-TauT antibody (PA5-37460, Invitrogen, 1:100 dilution). The slides were then detected using the HRP-conjugated compact polymer system and counterstained with hematoxylin. All the slides were observed with a light microscope and five fields were randomly selected for observation in each slide. For immunofluorescence, paraffin-embedded liver tissues were rehydrated and treated by heat mediated antigen retrieval using citrate buffer. The slides were incubated with donkey serum for 30 min at room temperature and then incubated with primary antibodies overnight at 4 °C. Anti-TauT antibody (PA5-37460, Invitrogen, 1:100 dilution), Anti-CK7 (abcam, ab181598, 1:8000 dilution) were used. Slides were then incubated with fluorochrome-conjugated secondary antibodies (1:500, Molecular Probes/Invitrogen) and observed with laser confocal microscopy. (Carl Zeiss, Jena, Germany).

### Quantitative PCR with reverse transcription (RT-qPCR)

Total RNA was extracted from RBE cells using RNAiso Reagent following the manufacturer’s instructions. Reverse transcription was performed on 1 μg of total RNA using the PrimeScript RT Reagent Kit at 37°C for 15 minutes. RT-qPCR was conducted on an Applied Biosystems StepOnePlus Real-Time PCR System (Thermo Fisher Scientific) using SYBR Premix Ex Taq II. The mRNA levels of target genes were quantified using the relative standard curve method and normalized to β-actin levels. Triplicate qPCR analyses were performed. Primer sequences were provided in Extended Data Table 3.

### Cell Senescence and Cell Proliferation assay

To assess the role of wild type TauT and mutations in regulating RBE cells anti-senescence and proliferation. Transfected WT or mutant TauT RBE cells were untreated or treated with 2μM DOX in media for 2hours to induce senescence. RBE cells were then treated with 1mM taurine for 24h. Trypsin cells, resuspended in 1X PBS, and fixed in 4% paraformaldehyde for 10 minutes at room temperature, then stained with the CellEventTM Senescence Green Probe (C10840, Invitrogen) for 120 minutes in a 37°C incubator with no CO2. Cells were washed in 1X PBS with 1% BSA, then resuspended in 1X PBS. Data was acquired on BD FACSClesta (BD Bioscience). A Cell-Light EdU Apollo488 In Vitro Kit (RioBio, C10310-3) was used for detecting cell proliferation assay, which cells were seeded onto 12-well culture plates and incubated with culture medium containing 20μM EdU solution for 2h after indicated treatment. According to the manufacturer’s protocol, cells were fixed by 4% formaldehyde for 30 minutes and incubated with 0.5% Triton X-100 for 10 minutes. Approximately 10^5^ cells were incubated with freshly prepared Apollo® staining reaction buffer in the dark for 30 min. Cells were washed in 0.5% Triton X-100 five times and then detected by LSDFortessa X20 (BD).

### bulk-RNA seq

We performed RNA-seq to determine the differentially expressed genes (DEGs) in DOX-treated RBE cells from WT and mutation TauT overexpression. Total RNA extracted from 2 groups (n=3 per group) was subjected to next-generation sequencing (NGS) to obtain deep coverage RNA-seq data. Transcriptome libraries were generated from the double-stranded cDNA, and further sequenced on an Illumina HiSeq instrument. Gene set enrichment analysis (GSEA) based on Gene ontology, KEGG and Reactome pathways was used to evaluate functional changes of each sample.

### Western blot

Cells or liver tissues were lysed with RIPA Lysis Buffer (Beyotime) for Western blot analyses. Lysates were centrifugated for 15min at 12000rpm, 4 ℃. Proteins were then fractionated by sodium dodecyl sulfate-polyacrylamide gel electrophoresis (SDS–PAGE) and transferred to PVDF membranes (Biorad). Membranes were blocked in 4% BSA solution for 1 h at room temperature. Antibodies against the TauT (PA5-37460, Invitrogen, 1:1000 dilution), p16 (Abcam, Ab108349, 1:1000 dilution), p21 (Abcam, Ab109520, 1:1000 dilution), ATF3 (CST, mAb #18665, 1:1000 dilution), HRP-conjugated β-Actin (Aksomics, KC-5A08, 1:5000 dilution), and HRP-conjugated secondary antibodies (Aksomics, KC-RB-035, 1:5000 dilution) were used. Immune complexes were detected using Immobilon^TM^ Western Chemiluminescent HRP Substrate (Millipore). The membranes were then visualized and quantified using an imaging system (Biorad).

### Statistical analysis

Data were shown as means ± standard error of the mean (SEM) and were analyzed using the GraphPad Prism 10. Statistical analysis was conducted using Student’s t test or Mann-Whitney U test between two groups and one way ANOVA in multiple groups. P<0.05 was considered statistically significant.

## Authors contributions

H.Z. initiated this project collaborated with N.N.C. H.Z. designed the expression constructs, purified the TauT protein samples, prepared cryo-EM grids, calculated cryo-EM data, built and refined structural models, prepared figures and wrote the initial manuscript. N.N.C. conducted the transport assay, cell proliferation, RT-PCR, western blot, RNA Seq, other aging related assay, prepared figures. H.E.X., in collaborations with X.M., supervised the project. H.E.X. modified the manuscript.

---

**Extended Data Fig. 1.**
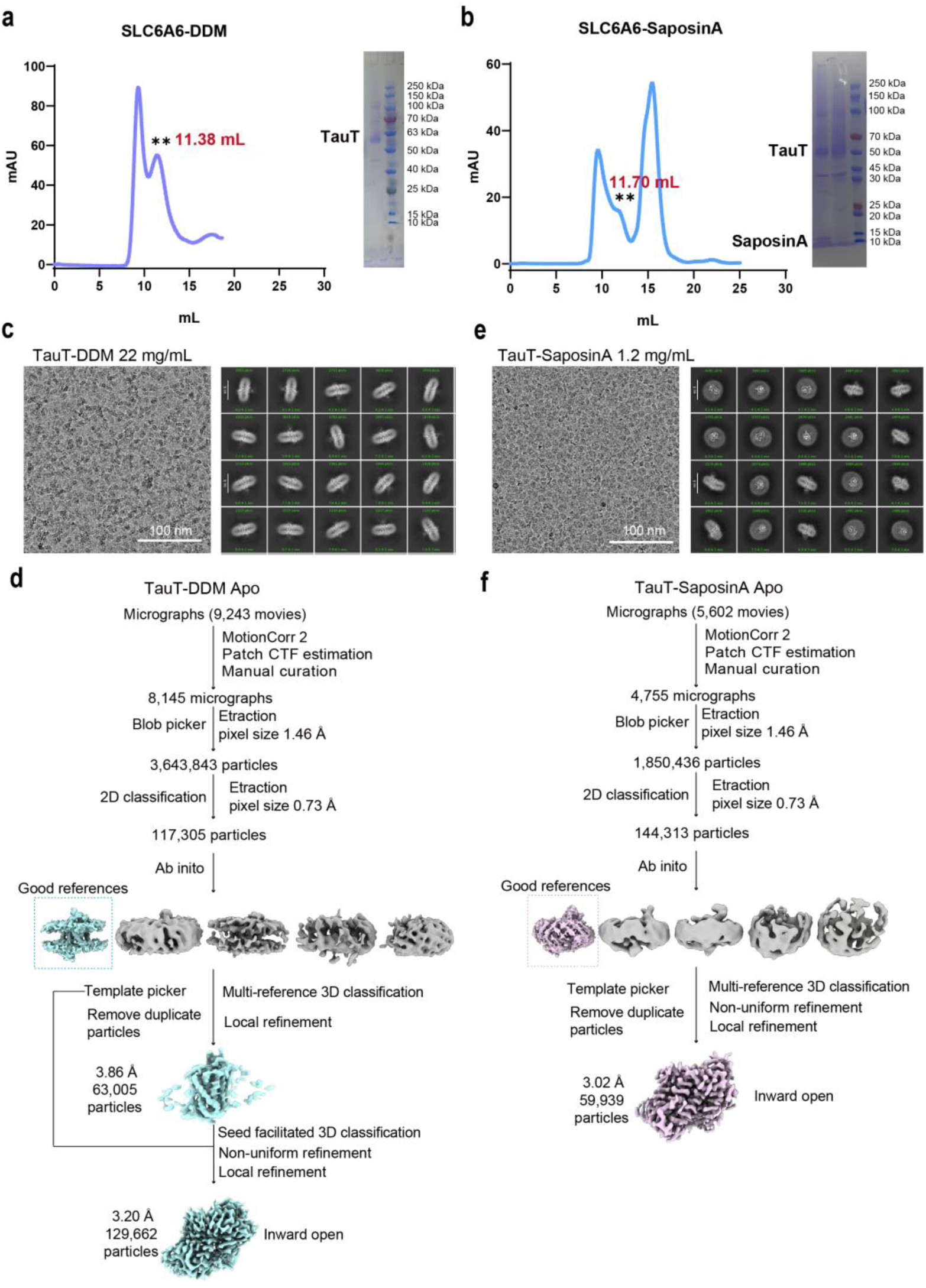
Cryo-EM structural determinations of human wild-type TauT at the apo state in the conditions of DDM and Saposin A nanodisc. **a, b,** The size-exclusion chromatography evaluations and the SDS-PAGE analysis results of the human TauT in the detergent of 0.012% DDM (**a**), and the TauT reconstituted into Saposin A Nanodisc (**b**). **c, e,** Representative cryo-EM micrographs and 2D classifications of TauT-DDM (**c**) and TauT-Saposin A (**e**), the scale bar located at the lower right in the micrographs. **d, f,** Data processing workflow of TauT-DDM (**d**) and TauT-Saposin A (**f**) by Cryosparc4.3.1 and ChimeraX.

**Extended Data Fig. 2.**
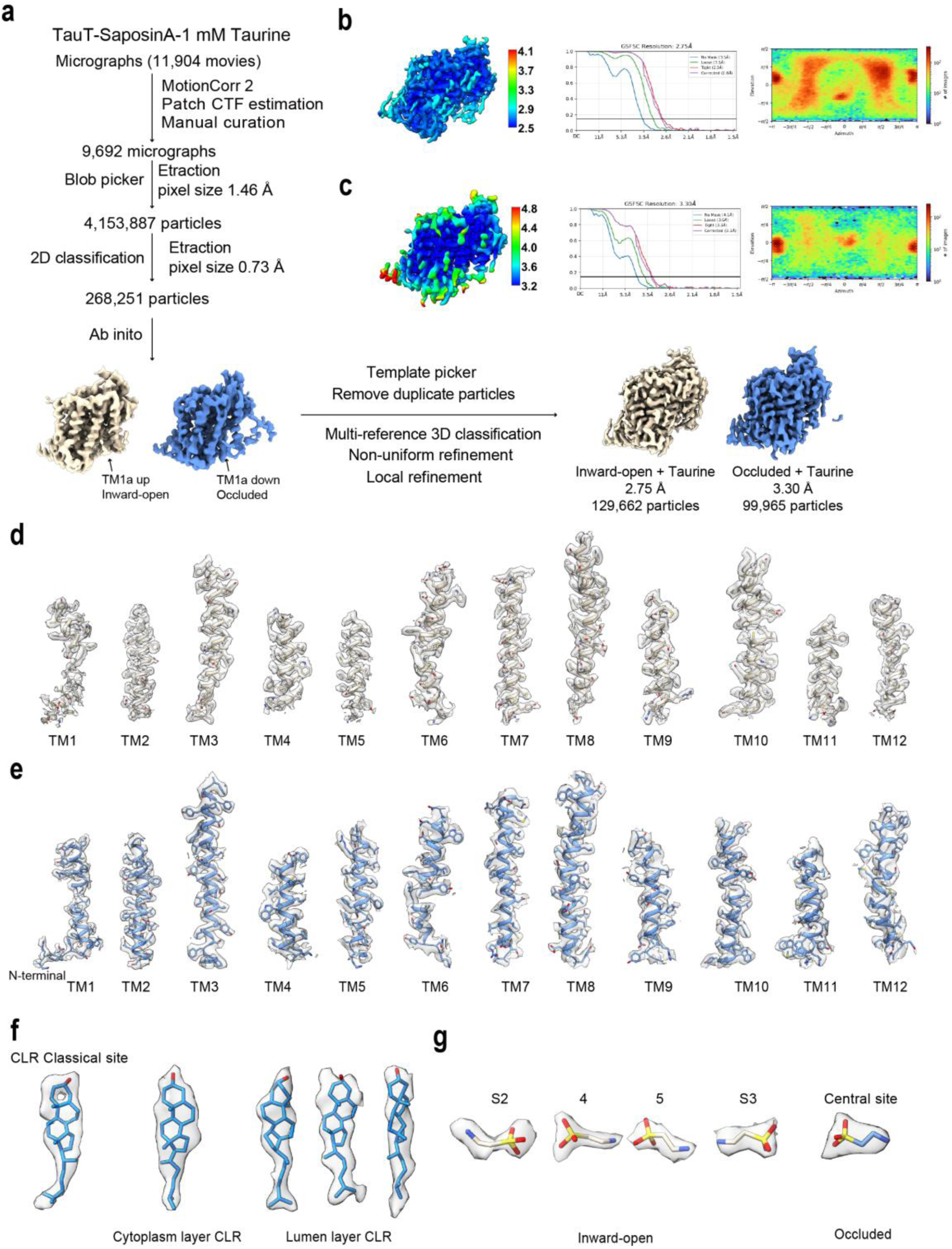
Cryo-EM analyses of TauT in taurine-bound states. **a,** Summary of image processing procedures of TauT in the presence of taurine. All procedures were done with cryoSPARC, except for motion correction which was done with RELION. **b, c,** Local resolution evaluations, Fourier shell correlation (FSC) curves and angular distributions of particles for the final 3D reconstructions of the inward-open (**b**) and occluded (**c**) conformations, respectively. **d, e,** Cryo-EM densities of the transmembrane helices of the inward-open (**d**) and occluded (**e**) conformations, respectively. **f,** Cryo-EM densities of the cholesterols in the taurine-bound TauT at the inward-open conformation. **g,** Cryo-EM densities of the taurine molecules in the taurine-bound TauT.

**Extended Data Fig. 3.**
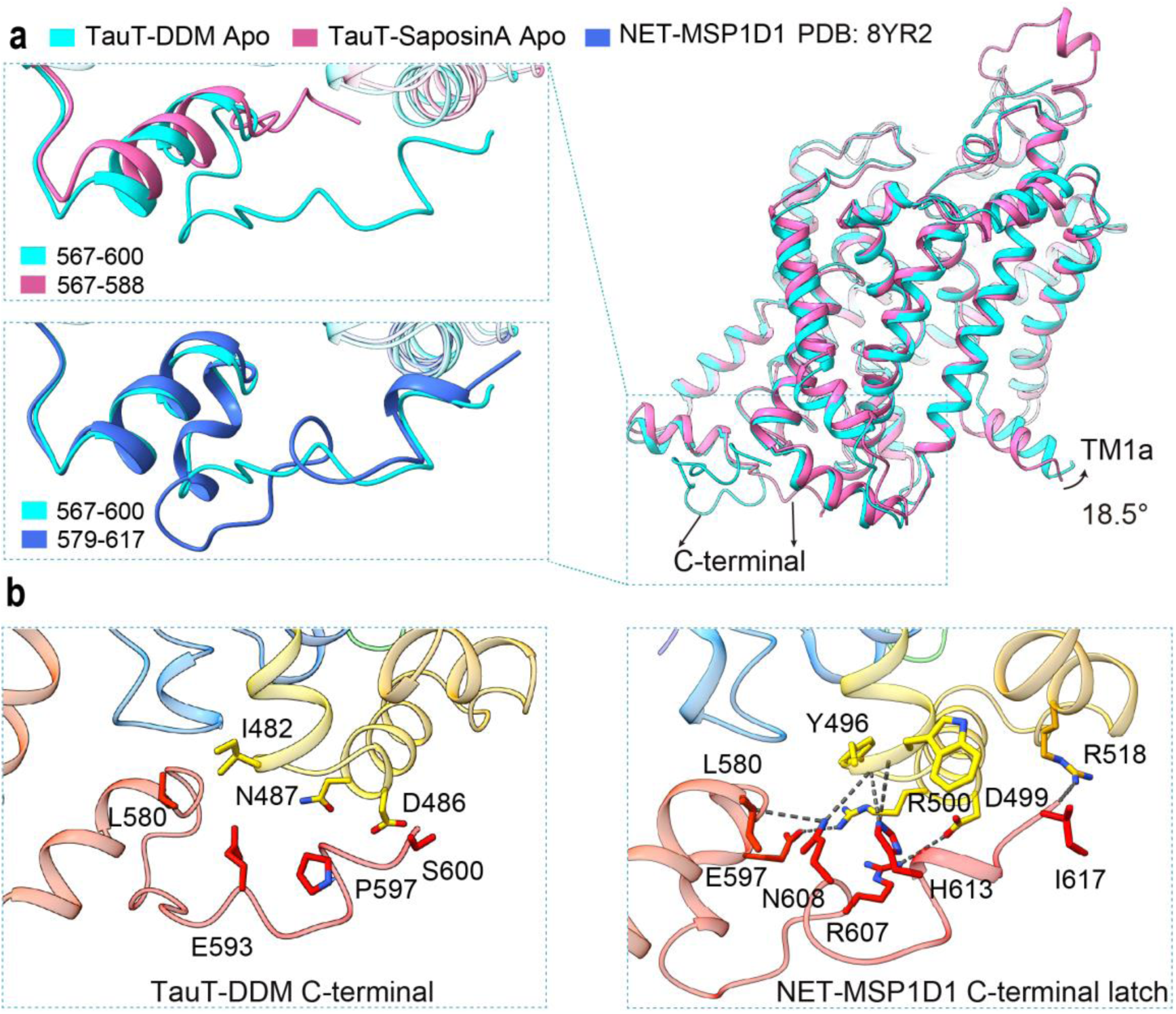
Structural comparisons of TauT in the DDM and Saposin A nanodisc. **a,** Structural comparisons of the overall and C-terminal structures of TauT-DDM and TauT-Saposin A in the apo state. **b,** Structural comparisons of the C-terminal structures of TauT-DDM and NET-MSP1D1 (PDB: 8YR2).

**Extended Data Fig. 4.**
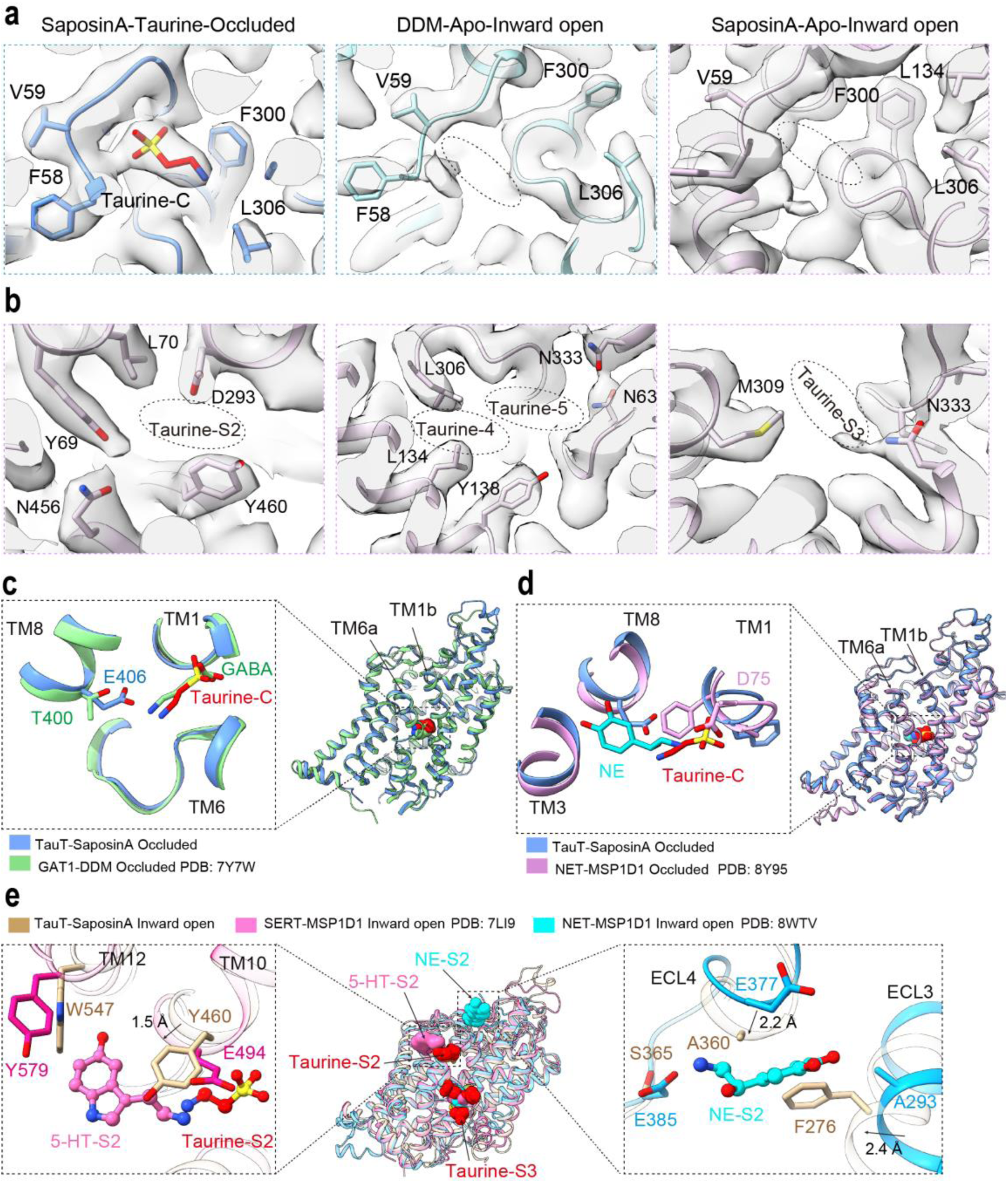
Structural analysis of the taurine binding sites. **a,** Cryo-EM densities (gray surface) corresponding to taurine at central binding site at taurine-bound occluded conformation (left), at TauT-DDM apo state (medium) and at TauT-Saposin A apo state (right), respectively. **b,** Cryo-EM densities (gray surface) corresponding to taurine binding site S2 (left), taurine-4, taurine-5 (medium) and taurine binding site S3 (right) at TauT-Saposin A apo state, respectively. **c,** Structural comparison of the taurine and GABA central binding pockets. **d,** Structural comparison of the taurine and NE central binding pockets. **e,** Structural comparison of the allosteric binding sites of taurine in TauT, 5-HT in SERT and NE in NET. Taurine, 5-HT and NE coloured in red, pink and cyan, represented as spheres, respectively.

**Extended Data Fig. 5.**
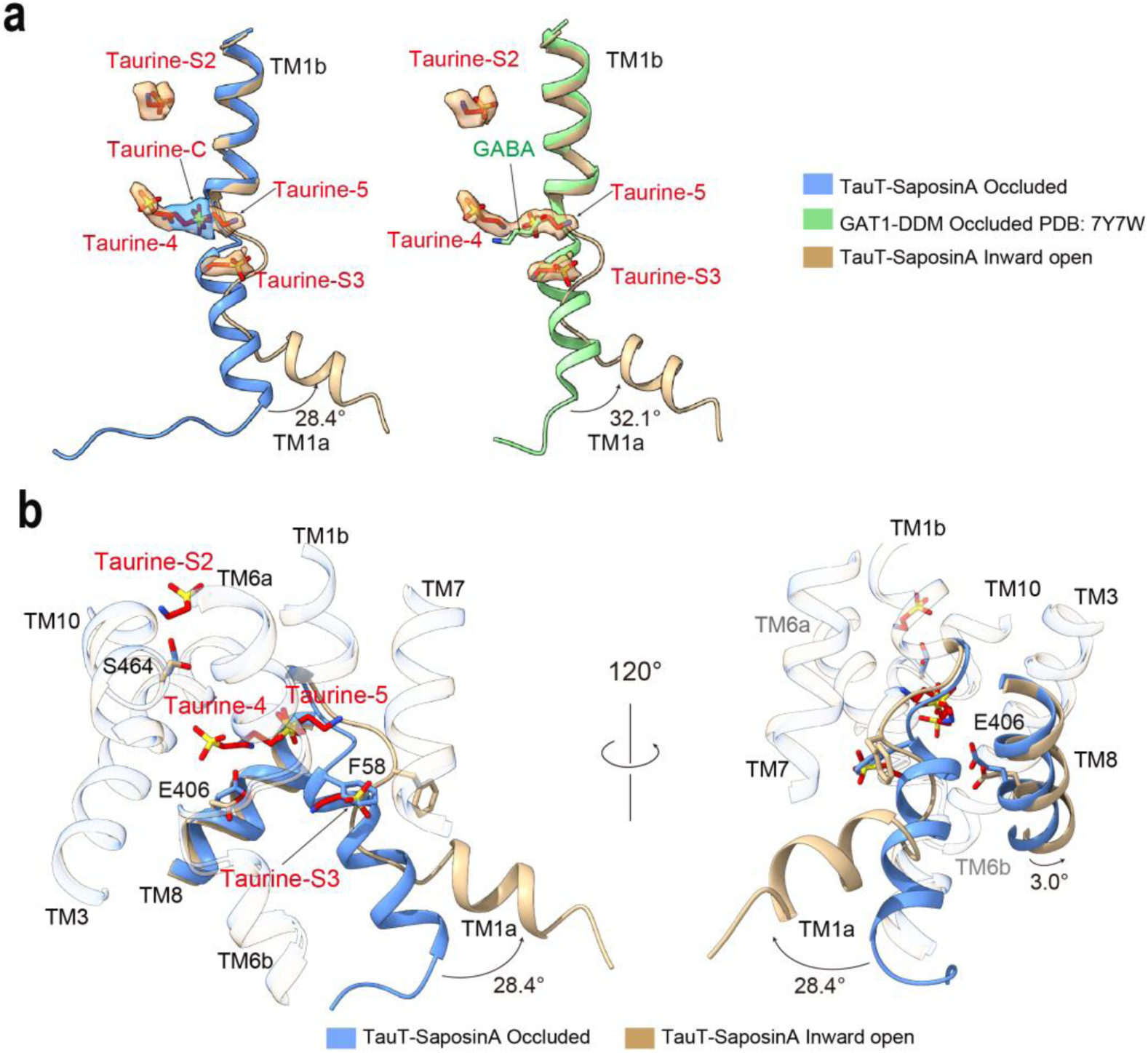
Structural analysis of occluded-inward transition. **a,** Structural comparisons of the TM1 and substrate binding sites of TauT and GAT1. **b,** Structural analysis of occluded-inward transition, especially focus on the TM1a, TM8 and taurine molecules.

**Extended Data Fig. 6.**
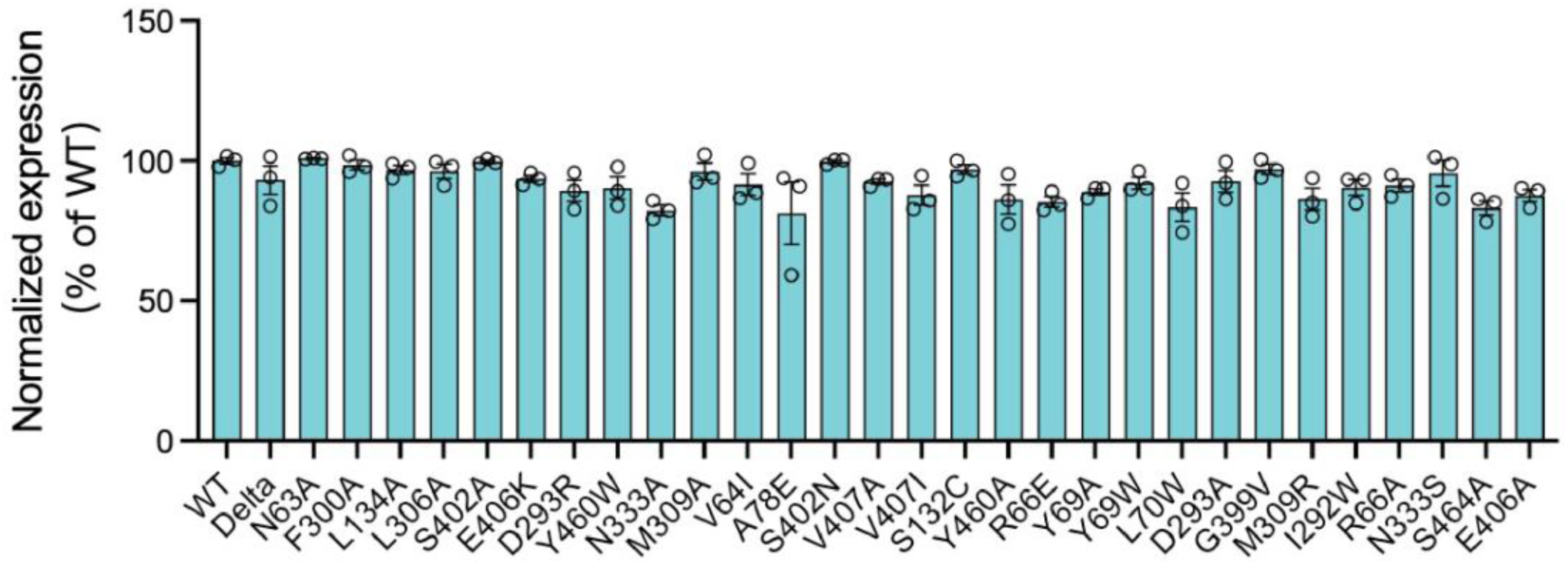
Quantification of surface expressions for wild-type TauT and its mutants was performed. Mutants of TauT expression levels were normalized to wild-type TauT using FACS. Data are presented as mean ± S.E.M., with n = 3 biologically independent experiments.

**Extended Data Fig. 7.**
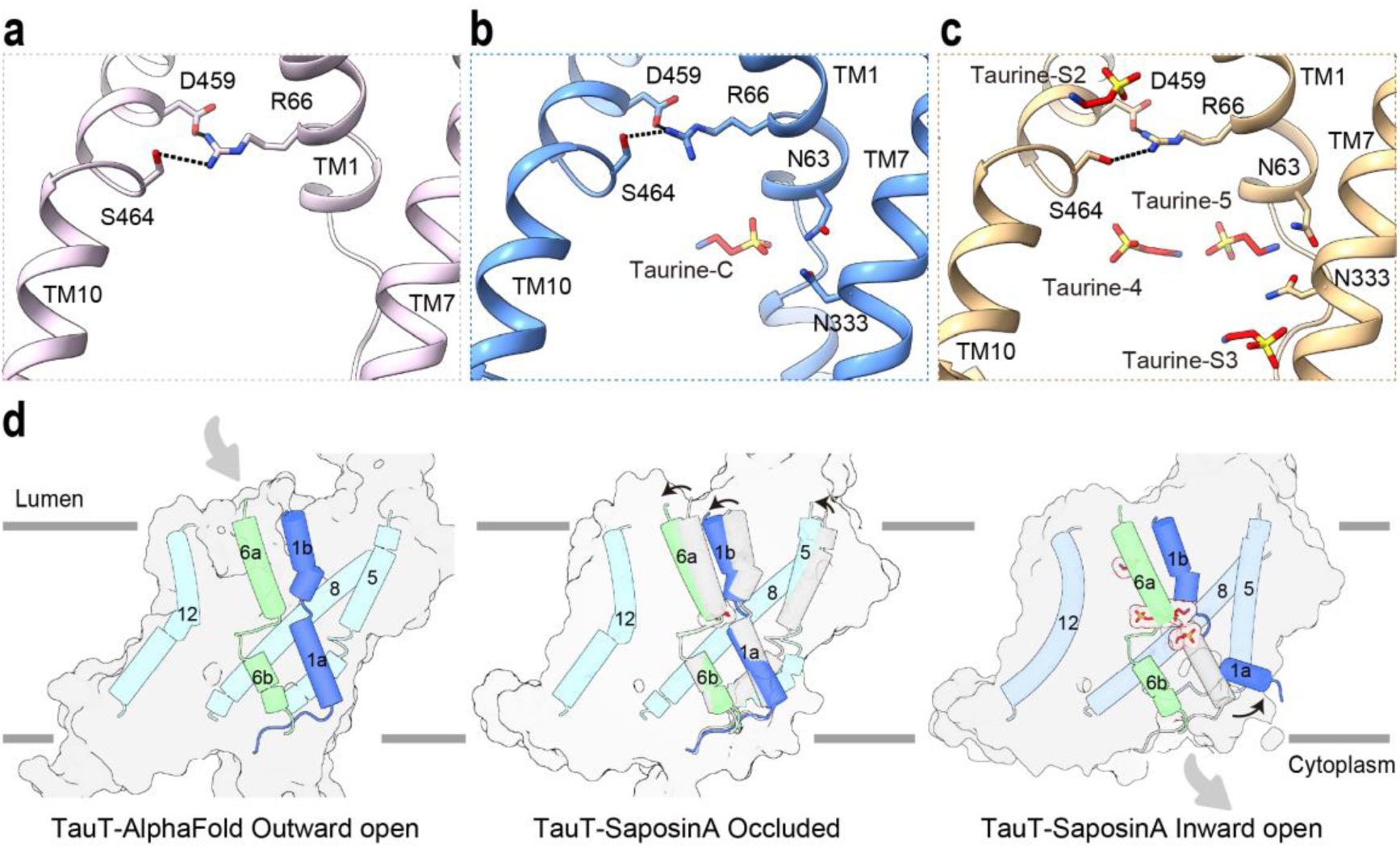
Outward entrance gate and transport mechanism of TauT. **a, b, c,** Structural details of the outward entrance gates formed by R66, S464 and D459 in the apo state (**a**), occluded conformation (**b**) and inward-open conformation (**c**) of TauT. **d,** Mapping of the transport mechanism of TauT. Key helices involved in the conformational changes are colored and labeled, the taurine molecules are colored in red and shown as surface. The outward open structure is predicted by AlphaFold2 (AF-P31641).

**Extended Data Fig. 8.**
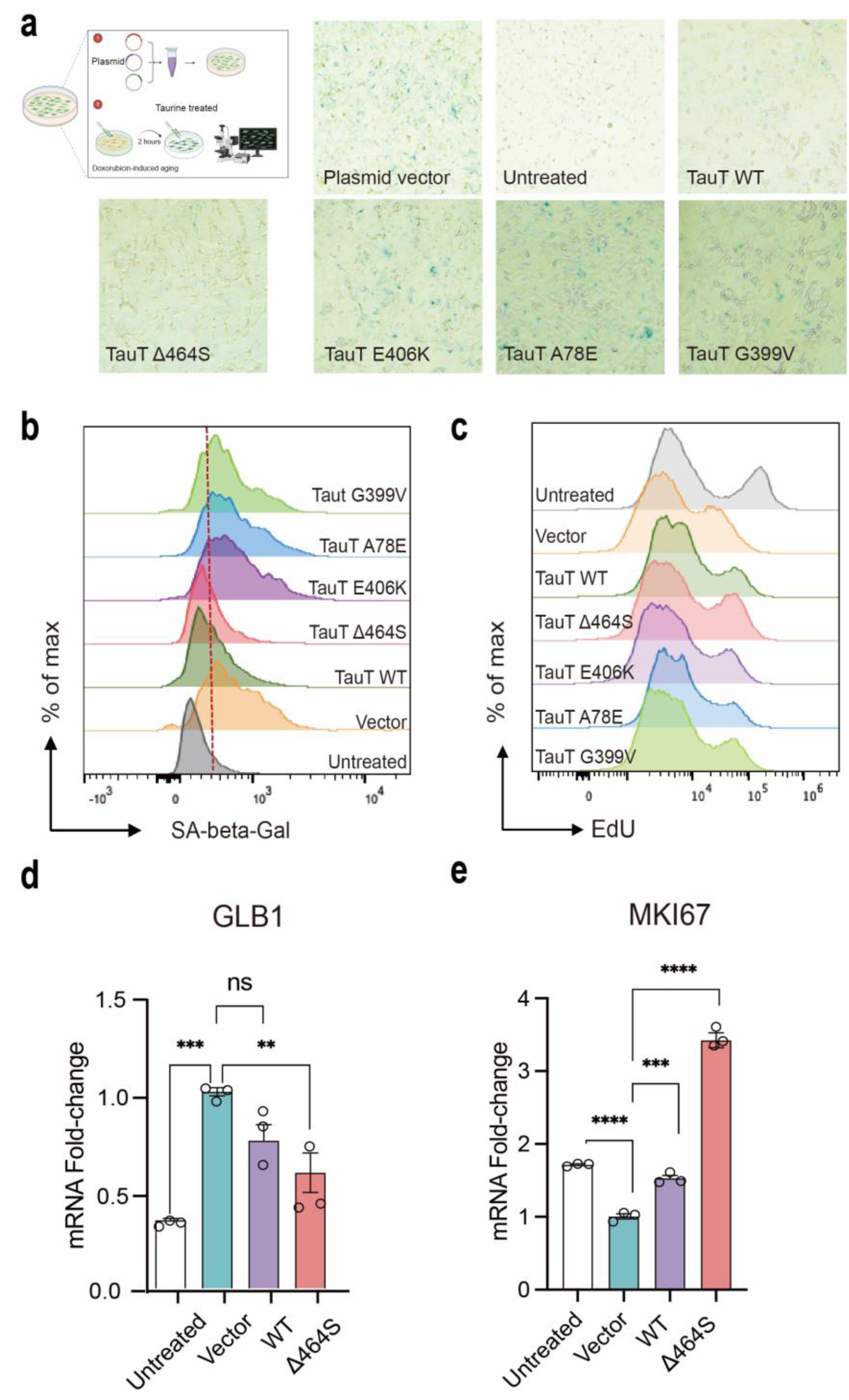
Augment taurine uptake alleviates the biliary epithelial cells aging. **a,** Senescence associated β-galactosidase (SA-β-Gal) staining (blue-stained cells) of RBE cells in doxorubicin treated and transfected with wilt-type TauT or TauT mutants. **b,d,** The representative plots of β-galactosidase expression (**b**) and GLB1 (**d**) of RBE cells in doxorubicin treated and transfected with wilt-type TauT or TauT mutants. **c,e,** The representative plots of EdU levels (**c**) and MKI67 expression (**e**) of RBE cells in doxorubicin treated and transfected with wilt-type TauT or TauT mutants.

**Extended Data Fig. 9.**
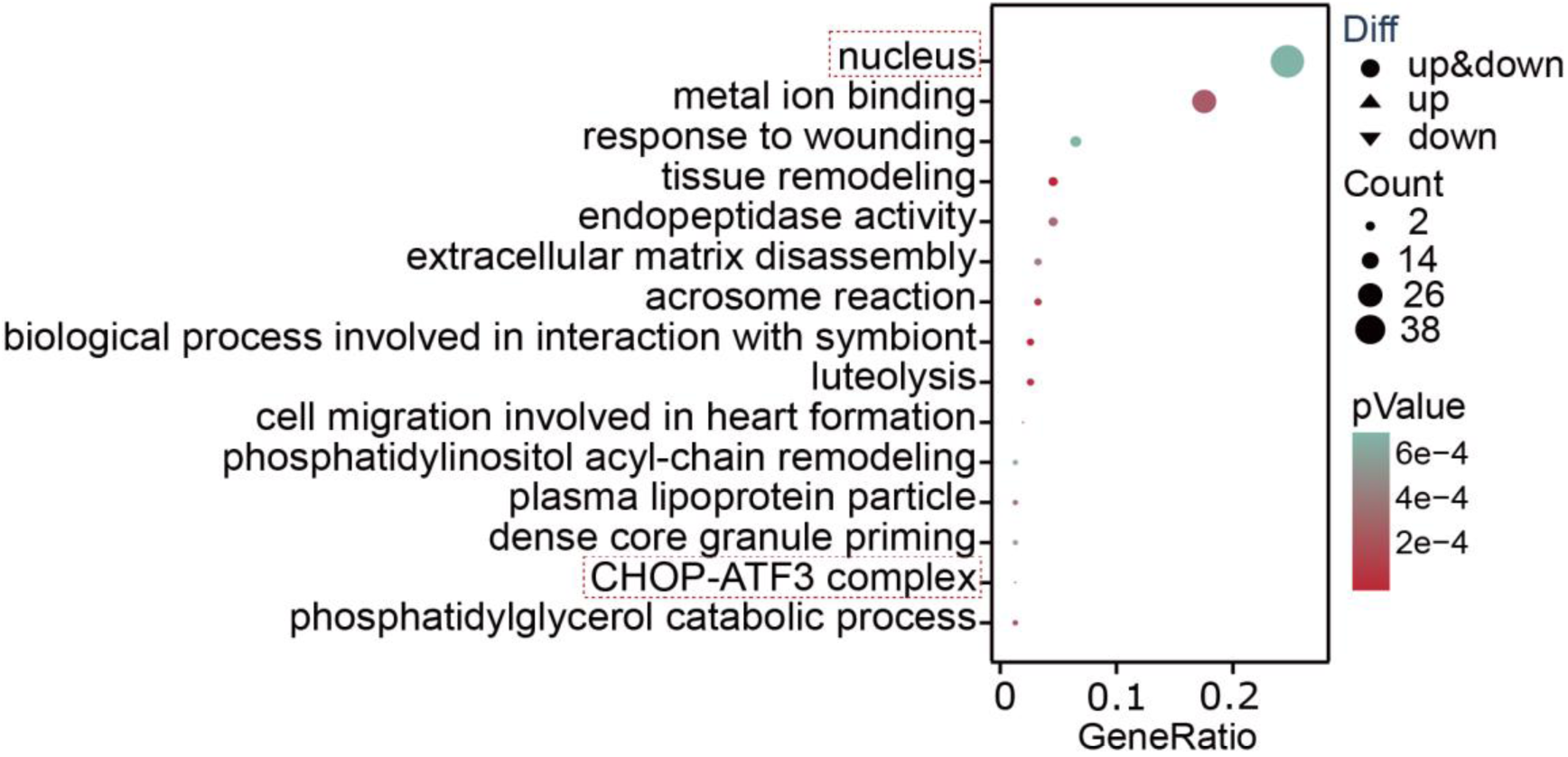
Transcriptome analysis of RBE cells in doxorubicin treated and transfected with wilt-type TauT.

**Extended Data Table1.**
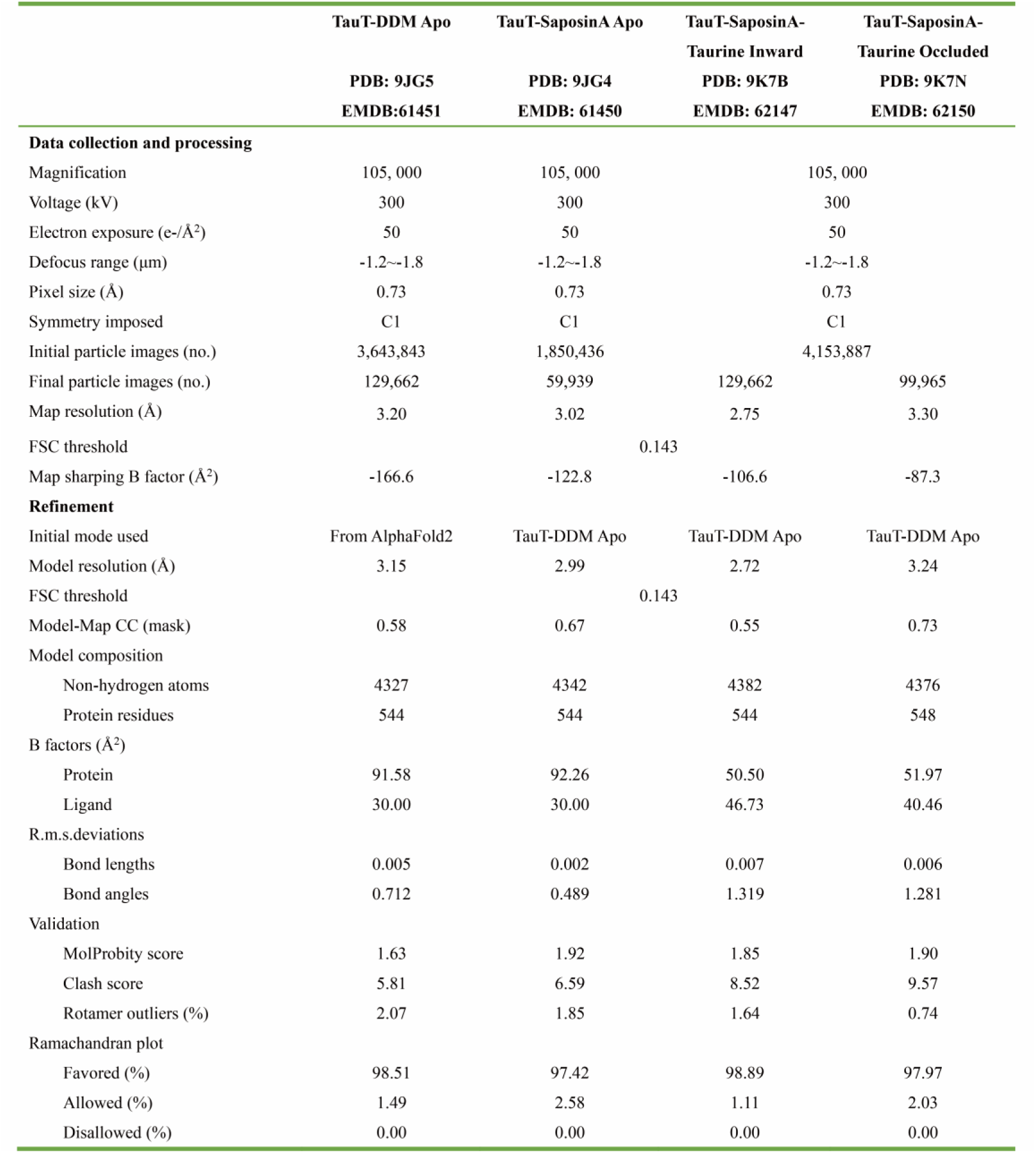
Cryo-EM data collection, refinement and validation statistics.

**Extended Data Table2.** Human subjects and samples used in the study.

| Sample | Condition | Aetiology of Liver Disease | Age | Gender | ALT(U/L) | AST (U/L) | ALP (U/L) | GGT (U/L) | TBIL(umol/L) |
| --- | --- | --- | --- | --- | --- | --- | --- | --- | --- |
| Healthy 1 | Healthy | - | 28 | F | 6 | 12 | 39 | 12 | 9.7 |
| Healthy 2 | Healthy | - | 41 | M | 19 | 18 | 43 | 31 | 11.1 |
| Healthy 3 | Healthy | - | 28 | F | 11 | 15 | 96 | 11 | 11.3 |
| Healthy 4 | Healthy | - | 36 | M | 19 | 17 | 95 | 17 | 21.8 |
| Healthy 5 | Healthy | - | 36 | F | 20 | 23 | 62 | 12 | 8.1 |
| Cirrhotic 1 | Cirrhotic | PBC | 44 | F | 63 | 135 | 140 | 88 | 426.7 |
| Cirrhotic 2 | Cirrhotic | PBC | 65 | M | 21 | 41 | 97 | 18 | 25.4 |
| Cirrhotic 3 | Cirrhotic | PBC | 48 | F | 45 | 100 | 293 | 90 | 135.6 |
| Cirrhotic 4 | Cirrhotic | PBC | 55 | F | 42 | 115 | 252 | 63 | 265.3 |
| Cirrhotic 5 | Cirrhotic | PBC | 41 | F | 128 | 122 | 336 | 357 | 187.7 |

**Extended Data Table3.**
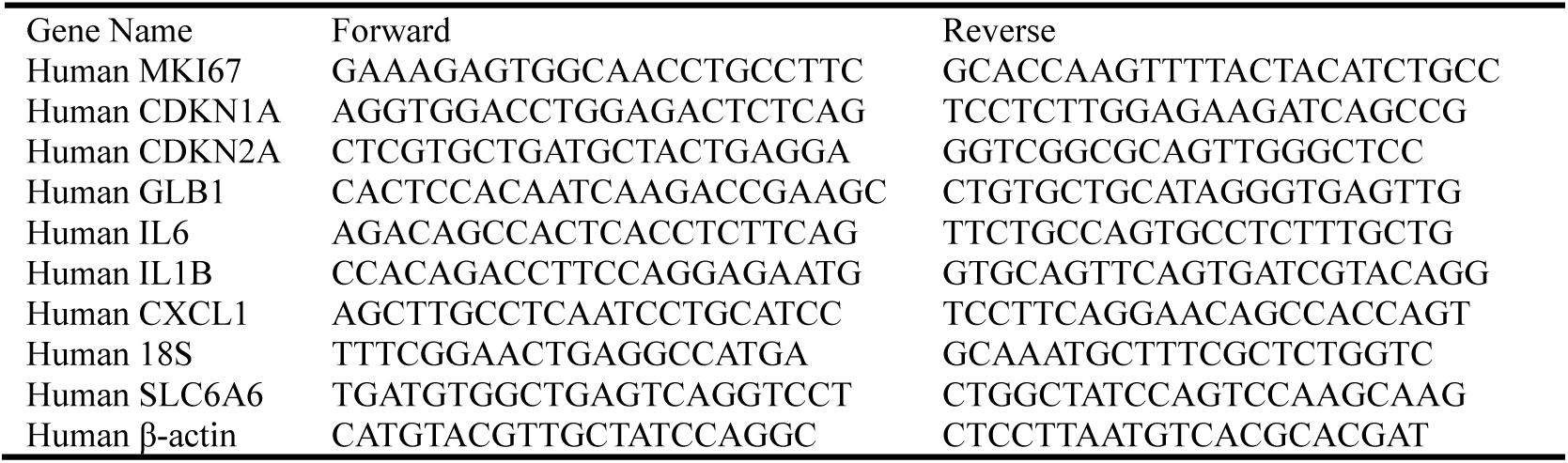
Primers used in the study.

